# dAMN: a genome scale neural-mechanistic hybrid model to predict bacterial growth dynamics

**DOI:** 10.64898/2026.03.04.709593

**Authors:** Jean-Loup Faulon, Danilo Dursoniah, Paul Ahavi, Antoine Raynal, Enrique Asin-Garcia

## Abstract

**Summary:** This study presents dAMN, a hybrid neural–mechanistic model that integrates neural networks with genome-scale dynamic flux balance analysis (dFBA) to predict bacterial growth curves across diverse nutrient environments. dAMN uses neural networks to infer dynamic behavior from initial metabolite concentrations, while mechanistic constraints ensure stoichiometric and thermodynamic consistency based on genome scale metabolic models. dAMN is trained on *E. coli* and *P. putida* experimental growth data from media containing various combinations of sugars, amino acids, and nucleobases, and evaluated on two test sets: one for forecasting over time and another for predicting growth dynamics on unseen media. dAMN achieved high predictive power (R^2^ ≥ 0.9), successfully reproducing growth and substrate depletion dynamics including acetate overflow and glucose-acetate consumption shift for *E. coli*. An interesting innovation of dAMN is the treatment of the lag phase, enabling realistic adaptation dynamics absent from standard dFBA models. dAMN stands out for its ability to generalize across combinatorial nutrient inputs and produce full growth-curve predictions from minimal input data.

**Availability and implementation:** The dAMN software, along with the associated models and data, is available at https://github.com/brsynth/dAMN-main-release and via DOI 10.5281/zenodo.17908125

## Introduction

A central challenge in modeling microbial metabolism is predicting growth dynamics across diverse nutrient environments. Most of the mechanistic and hybrid approaches struggle to generalize across media and rarely capture lag-phase adaptation. Dynamic Artificial Metabolic Networks (dAMN) address this gap. By embedding neural network components within a mechanistic dynamic flux balance analysis (dFBA) framework, dAMN learns substrate-dependent flux adjustments and lag-phase delays directly from nutrient environments while preserving the stoichiometric constraints of FBA. This enables accurate prediction of full growth curves including lag-phases across diverse media conditions.

Mathematical modeling of cellular metabolism has traditionally relied on kinetic models or constraint-based approaches. While kinetic models provide mechanistic accuracy, they require extensive parameterization and are not scalable to genome-scale networks. In contrast, Flux Balance Analysis (FBA) offers a scalable, stoichiometry-based framework for computing optimal steady-state flux distributions without requiring kinetic parameters (Orth *et al*., 2010). However, its steady-state assumption precludes direct simulation of batch or fed-batch processes, where extracellular conditions and intracellular fluxes change over time.

Dynamic Flux Balance Analysis (dFBA) extends FBA to temporal processes. Originally outlined by Varma and Palsson (Varma and Palsson, 1994) and formalized by Mahadevan *et al*. (Mahadevan *et al*., 2002) dFBA iteratively solves a sequence of FBA problems alongside integration of extracellular metabolite concentrations. Although dFBA captures time-varying behaviors, it often produces biologically unrealistic instantaneous shifts in metabolic fluxes and faces numerical stiffness.

To address these issues, several extensions have been proposed. Enzyme-constrained FBA (ecFBA) limits reaction rates based on enzyme capacity (Sánchez *et al*., 2017), while enzyme-change constrained models (Karlsen *et al*., 2023) explicitly bound the rate at which proteome fractions reallocate. Such models better reproduce phenomena like diauxic lags in *E. coli*, however these models require additional data beyond media composition. Other efforts extend dFBA to microbial consortia (Stolyar *et al*., 2007)(Tzamali *et al*., 2011) or combine it with statistical surrogates (Negahban *et al*., 2025).

Recently, hybrid methods that integrate data-driven machine learning with mechanistic modeling have emerged as alternatives to purely kinetic or constraint-based approaches. Physics-Informed Neural Networks (PINNs) embed ODE systems directly in neural architectures, enabling estimation of hidden dynamics and parameters from sparse data (Raissi *et al*., 2019) and (Yazdani *et al*., 2020). PINNs require an explicit ODE model for growth that is rarely available at the genome scale. Neural-ODE based metabolic models (Rathod *et al*., 2024) are genome scale, they infer latent dynamic states from gene expression and decode them into fluxes but cannot reproduce growth curves. Artificial Metabolic Networks (AMNs), also genome scale, embed neural networks inside the FBA framework to improve flux prediction while reducing data requirements (Faure *et al*., 2023), however, AMNs do not handle metabolite or biomass concentrations. Other neural surrogates have been proposed, among theses: (Song *et al*., 2025) trains ANNs as FBA surrogates for Shewanella oneidensis in reactive-transport models, Dehkordi *et al*. (Dehkordi *et al*., 2023) uses convolutional networks to simulate Saccharomyces cerevisiae growth for fast control, COSMIC-dFBA (Gopalakrishnan *et al*., 2024) couples a data-driven cell-state classifier with dFBA to predict time-resolved metabolite profiles, and (Negahban *et al*., 2025) constrain dFBA with partial least squares regression on kinetic parameters. However, these methods typically operate on fixed or narrow sets of substrates, treat the machine learning and dFBA components separately, or do not explicitly simulate extracellular metabolite depletion and growth trajectories, so they cannot systematically predict lag phases or generalize growth behavior across diverse media compositions.

To palliate these shortcomings, we present dynamic Artificial Metabolic Networks (dAMN), a hybrid model combining neural networks with mechanistic constraints from dFBA. Trained on experimental growth curves across combinatorial nutrient mixtures, dAMN predicts reaction fluxes and lag-phase dynamics from initial conditions and integrates these to produce full growth trajectories. We detail the model architecture, training procedure, and predictive performance in the following sections. Additional methods and results can be found in the Supplementary Information (SI), and links to the SI are provided whenever necessary.

## Method

The methodological approach presented here is structured into three distinct subsections. The first subsection describes the dAMN model, detailing its hybrid neural-mechanistic architecture, key equations, and loss functions used for training and predictions. The second subsection summarizes the experimental procedures used to generate the training datasets, including bacterial culture preparation and growth monitoring. The final subsection outlines the construction and use distinct training and test datasets for model validation.

### dAMN model

The overall architecture of the dAMN model is illustrated in Fig. S1 in SI. At its core, dAMN is a hybrid neural-mechanistic model integrating neural networks with dFBA. The neural network components 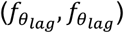 is designed to predict lag phase parameters 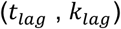 and reaction fluxes (*V*(*t*)), from medium composition (*C*_*true*_(0)). The mechanistic component applies stoichiometric and kinetic constraints to these flux predictions, ensuring biological plausibility.

The model can be formalized through the following equations:

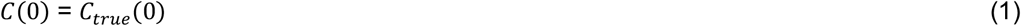

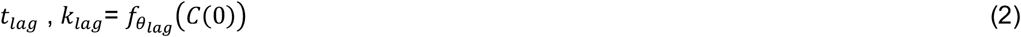

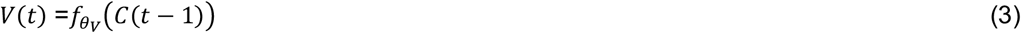

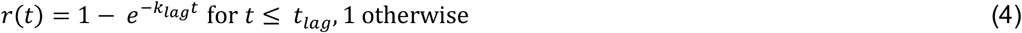

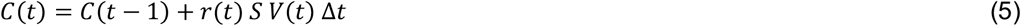

where:

*t* : dis Crete time in [0, *t*_*max*_] with Constan *t* spacing by Δ#

*C*_*true*_(*t*): measured Concentration at time *t* for all reference metabolites (biomass included)

*C* (*t*) : learned Concentration at time *t* for all metabolites

*V*(*t*) : learned flux at time *t* for all reactions

*r*(*t*) : a exponential function *r*(*t*) = 0 a*t t* = 0 and *r*(*t*) = 1 for *t* ≥ *t*_*lag*_

*t*_*lag*_ : learned lag *t*ime

*k*_*lag*_ : learned stiffness of the exponential function *r*(*t*)

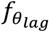 : feedforward neural ne*t*work with parame*t*ers θ_*lag*_

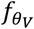 : feedforward neural ne*t*work with parame*t*ers θ_*V*_

*S* : stoichiometric matrix

Let us note that for internal metabolites *C*(*t*) = *C* (*t* ™ 1) as = *SV* (*t*) = 0 consequently Eq. (5) can be written as:

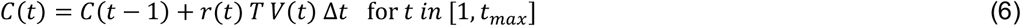

where *T*, derived from the stoichiometric matrix *S*, can be viewed as a transport matrix mapping reactions to external metabolites, *T*=-1 (+1) for inward (outward) reactions.

As in PINN we train our dAMN model in order to match both the observation (measured concentrations) and mechanistic constraints (dFBA constraints). The loss function given below reflect this multi-objective.

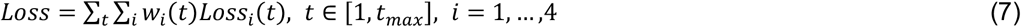

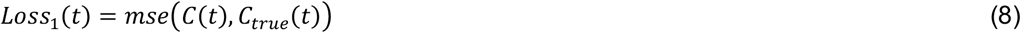

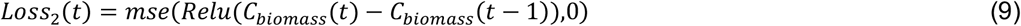

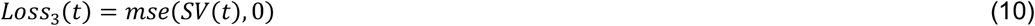

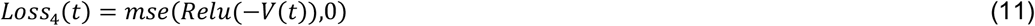

The loss function enforces predicted concentrations match measured concentrations (*Loss*_1_), biomass concentrations cannot reduce overtime (*Loss*_2_), the stoichiometric constraint is verified (*Loss*_3_), and all reactions are irreversible and therefore all fluxes are positive (*Loss*_4_). The weights, *w*_*i*_(*t*), are specific to each loss term and evolve overtime. Several functions, linear, exponential, cosines and sigmoid have been tested. Best results were obtained with the following exponential decay function:

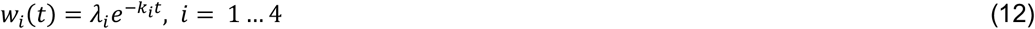

To determine the optimal values for the loss weighting (λ_*i*_) and decay parameters (*k*_*i*_) in the loss function, we performed hyperparameter optimization (see Results section).

We tested alternative architectures, in particular to model the lag-phase using a Hill function as in (Flassig *et al*., 2016), and to remove time decay in the loss function. These are described in SI Section 1 (Figs. S1, S2 and S3). In general, these alternatives do not outperform the architecture described by Eqs. (1–12).

#### Experimental datasets

We acquired three experimental datasets. The first one (referred as M28) is composed of *E. coli* MG1655 growth curves acquired on 280 different culture media. The second (named putida) is composed of *P. putida* KT2440 growth curves acquired on 81 different culture media. In both cases the strains were grown on an M9 minimal medium supplemented with combinations of sugars (D-glucose, D-xylose, succinate), 20 amino acids, and 5 nucleobases. Further details on experimental procedures are provided in SI section 2 for *E. coli* and SI section 3 for *P. putida*. The third dataset (referred as Millard) was extracted from (Millard *et al*., 2021). This dataset comprised concentration measurements at various timepoints for glucose, extracellular acetate and biomass for *E. coli* MG1655 grown on an M9 minimal medium supplemented with glucose and acetate. Additional information for that dataset is provided in SI section 4.

#### Training and test sets

Two training and test sets were generated for both *E. coli* (M28) and *P. putida*. In the first set, the forecast set, dAMN was trained using two-thirds of the growth time points from all available data, and predictions were made on the remaining one-third. In the second set, the media set, dAMN was trained on two-thirds of the available media using all time points, and predictions were made for media conditions not included in the training set. In addition, two training/testing datasets were generated using the *E. coli* Millard data mentioned above. In the first case, the training set was composed of measured concentrations for extracellular acetate and biomass (with glucose concentrations in test set), and in the second case the training set was composed of measured concentrations for glucose and biomass (with acetate concentrations in test set). Additional information for these datasets are provided in SI section 4.

## Results

Prior evaluating dAMN we needed to fine tune its architecture and hyperparameters. A critical dAMN parameter is the stoichiometric matrix used to build the mechanistic constraints. As we tested dAMN with *E. coli* MG1655 the stoichiometric matrix was obtained from *E. coli* genome-scale metabolic model iML1515 (Monk *et al*., 2017). This metabolic model comprises 1877 metabolites, 2712 reactions, and 1516 genes. To ensure all reactions were irreversible, reversible reactions were duplicated (one forward, one reverse) following the approach described in (Faure *et al*., 2023), resulting in 3682 irreversible reactions and a stoichiometric matrix of dimensions [1877, 3682]. To construct the transport matrix *T*, we extracted rows and columns from the stoichiometric matrix corresponding to external metabolites, resulting in a transport matrix of dimensions [29, 3682]. The 29 matrix rows included the 28 medium metabolites and the biomass for which concentration was monitored. The ‘*reversible reaction duplicated*’ iML1515 model was used for the *E. coli* M28 and Millard datasets mentioned earlier. For the Millard dataset extracellular acetate was added to the transport matrix. The same procedure was applied to *P. putida* model iJN1463 resulting in a stoichiometric matrix of dimensions [2153, 4034] and a transport matrix of dimensions [29, 4034].

We also searched the optimal loss weighting (*λ*_i_) and decay parameters (*k*_i_) of Eq. 12, performing hyperparameter optimization. To that end, a coarse random search was conducted over log-uniform ranges for *λ*_i_ ([0.001, 0.01, 0.1, 1.0]) and linear for *k*_i_ ([0.0, 0.25, 0.5, 0.75, 1.0]), training each model for 1000 epochs with early stopping. Each candidate was evaluated using the medium set (3-fold cross validation), and the average regression coefficient (R^2^) between predicted and experimentally measured growth on the test set served as the metric for model selection. The 10 best-performing parameter sets achieving the highest average and median test R^2^ are summarized in SI Table S1. The top optimal parameter set was subsequently used for further analyses. All experiments were conducted using fixed random seeds and consistent data splits to ensure reproducibility.

dAMN was first evaluated using the *E. coli* M28 and *P. putida* datasets mentioned earlier. In both cases, the forecast set and the media set were used to assess dAMN capacities in forecasting metabolite concentrations over time and predicting bacterial growth on unseen media conditions. Results are presented for *E. coli* M28 dataset in Figure 1 and in SI Fig. S6 for *P. putida*. Figure 1 (panels (a) and (b)) shows that the concentrations of substrates (glucose and succinate) follow expected biological behaviors, demonstrating gradual depletion corresponding to biomass accumulation. The forecast set results indicate strong predictive capability, exemplified by the close match between predicted and experimentally measured growth curves (panel (c)), with a median R^2^ of approximately 0.98 (panel (d)). The media set, which evaluates dAMN’s generalization to unseen media conditions, also exhibits robust predictions, as illustrated by the representative growth curve (panel (e)) and a median R^2^ value of around 0.96 (panel (f)).

**Figure 1.**
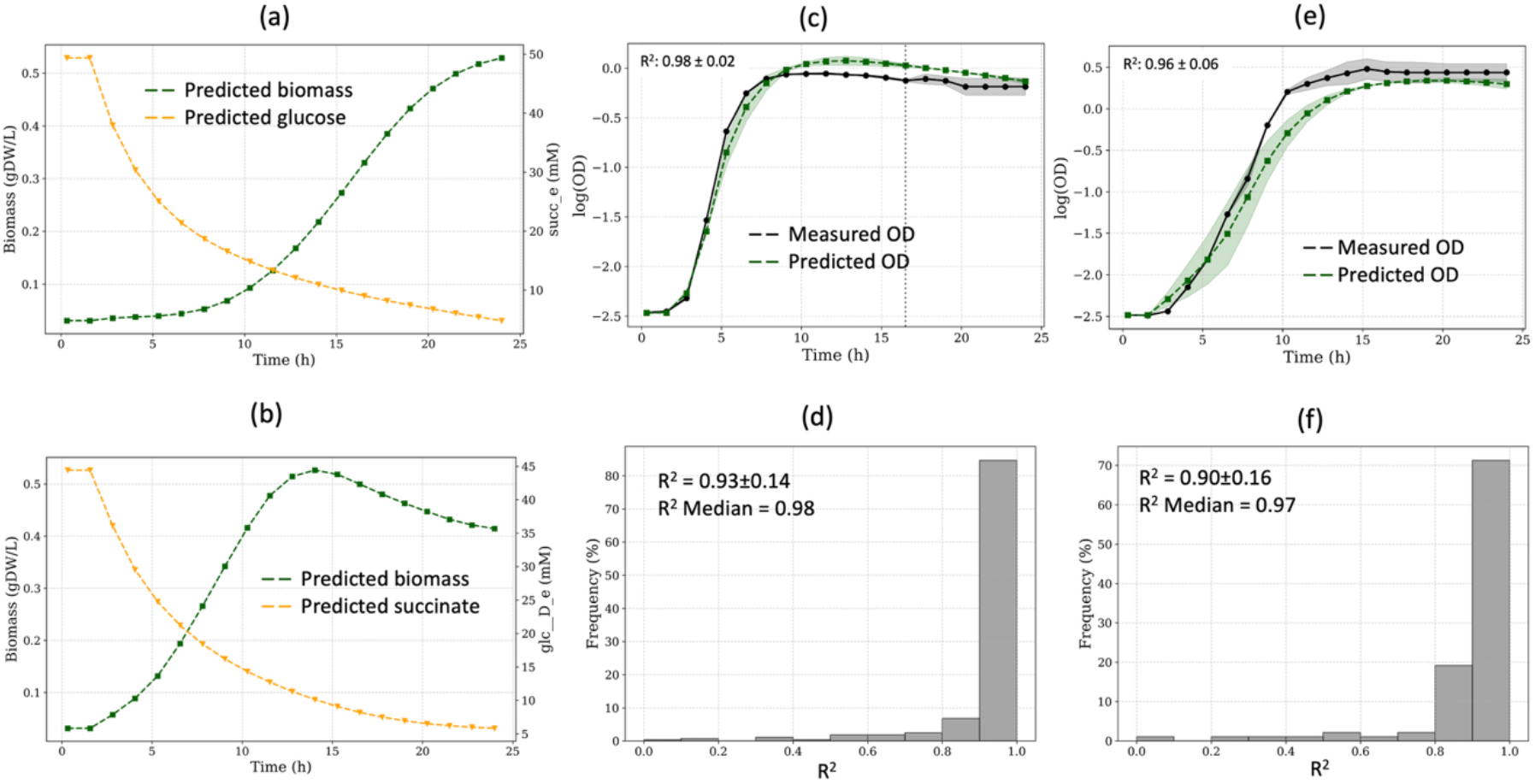
dAMN results for *E. coli* M28 dataset. (a) Predicted concentrations for biomass and D-glucose when the strain is grown on M9 with D-glucose only (medium number 1). (b) Predicted concentrations for biomass and succinate when the strain is grown on M9 with succinate only (medium number 156). (c) Example of a predicted growth curve compared to measured growth curve using the forecast training set (dashed line indicates when forecast begins). dAMN is trained on the first two-thirds of the datapoints, with predictions made for the final third. (d) Distribution of R^2^ values between measured and predicted growth curves for the forecast training set. (e) Example of a predicted growth curve compared to measured growth curve using the media training set. dAMN is trained on the first two-thirds of all media, and predictions are made for media not used during training. Displayed curve represents prediction for a medium not present in the training set. (f) Distribution of R^2^ values between measured and predicted growth curves for the media training set. Standard deviations (shaded areas in panels c and e) were obtained using three replicates for the experimental data, and running three models trained on the forecast and medium set using the top parameter set of Table S1.

As shown in SI Table S2, dAMN performance exhibits a moderate decline in predictive accuracy as the amount of training data was reduced (dropping from a mean R^2^ = 0.90 with 187 media in the training set to a mean R^2^ = 0.85 with only 93 media in training). Poorly performing examples (*cf*. Fig. S4 in SI) are mostly due to undetectable lag phases or sharp biomass drops during the stationary phase. Yet, dAMN appears to handle well different lag-phase lengths (Fig. S5 in SI).

These results collectively validate the model’s predictive accuracy and its generalization capacity to various nutritional contexts.

Results similar to those presented in Figure 1, were obtained with the *P. putida* dataset for both forecasting (*cf*. Fig. S6 panel (c) in SI where mean R^2^ = 0.94, median R^2^ = 0.98) and predicting growth curves of unseen media (*cf*. Fig. S6 panel (d) in SI where mean R^2^ = 0.90, median R^2^ = 0.94).

Because dAMN is able to predict substrate consumption concurrent with biomass increase (*cf*. Figure 1 panels (a) and (b)), we sought to assess this emergent behavior quantitatively. To this end, we used the Millard dataset and trained dAMN on biomass and extracellular acetate concentrations. Fig. S7 panel (a) in SI shows that glucose concentrations are accurately reconstructed under these conditions. Likewise, dAMN accurately predicts acetate overflow when trained on glucose and biomass concentrations (Fig. S7 panel (b) in SI). We compared these results (section 4 in SI) with those obtained with a PINN based on the ODE model of Millard (Millard *et al*., 2021). As is customary with PINNs, our PINN was trained both to fit the measured concentrations and to infer values for unknown kinetic parameters (Table S3 in SI). Fig. S7, panels (c) and (d), shows that dAMN outperforms PINN unless precise and accurate values for the unknown kinetic parameters are provided (Fig. S7, panels (e) and (f)).

To further investigate substrate consumption hierarchies, we trained dAMN exclusively on biomass and glucose time series, without providing acetate measurements during training, and extended the predictions beyond the experimental time window. In these forward simulations, dAMN predicted acetate consumption following glucose depletion, accompanied by continued biomass increase, consistent with classical diauxic behavior (Fig. S8 in SI), even though these phases were not present in the training data. These results indicate that dAMN can infer and reproduce multi-substrate utilization patterns from limited supervision.

## Discussion and conclusion

This study introduces dAMN, a hybrid neural–mechanistic model that integrates data-driven neural networks with mechanistic constraints from genome-scale metabolic networks to predict the full dynamics of microbial growth curves. The model was evaluated using two distinct test sets: a forecast set to assess temporal extrapolation, and a media set to evaluate its capacity to generalize to entirely unseen nutritional compositions. In both cases, dAMN demonstrated high predictive accuracy, achieving average R^2^ values ≥ 0.90 for both forecast and media sets acquired for *E. coli* (Figure 1) and *P. putida* (Fig. S6 in SI). These performances confirm that the model not only learns realistic time dynamics but also generalizes robustly across diverse and previously unobserved environmental conditions.

As with AMNs relative to FBA (Faure *et al*., 2023) a key advantage of dAMN over standard dFBA is its ability to infer uptake fluxes directly from extracellular medium concentrations, rather than requiring them to be measured or specified a priori. This data-driven estimation of uptake improves the accuracy of growth predictions, as illustrated by the dAMN–dFBA comparison carried out in Section 5 and Fig. S9 in the SI.

A particularly noteworthy outcome of our results is the predicted depletion pattern of substrates like glucose and succinate, as shown in Figure 1 panels (b) and (c). These results were obtained despite the fact that only the initial substrate concentrations were provided and no intermediate data was used during training. The model was nevertheless able to produce realistic substrate consumption curves that mirror biomass accumulation. This emergent behavior arises solely from the enforcement of the stoichiometric constraint =%(#) = 0, which ensures internal metabolite balance. Since substrate consumption is the only mechanism available to fuel biomass production, the model was constrained to infer realistic usage rates, an interesting outcome given the lack of direct supervision on substrate dynamics. This observation was quantitively validated with the *E. coli* Millard dataset where glucose consumption and acetated overflow could accurately be predicted when compared to measurements (section 4 and Fig. S7 in SI). Additionally, with forward simulations, dAMN was able to predict acetate consumption following glucose depletion (Fig. S8 in SI), even though this diauxic shift was absent from the training data.

The explicit modeling of the lag phase via exponential or Hill functions is another feature distinguishing dAMN. Classical dFBA implementations typically neglect this initial adaptation delay (*cf*. Fig. S9 in SI). While multiphase models based on empirical phase divisions (Moimenta *et al*., 2025) or enzyme-change constraints (Sánchez *et al*., 2017) and (Karlsen *et al*., 2023) have addressed lag phases, they often lack fully data-driven parametrization. Approaches incorporating kinetic delays or dynamic objectives (Waldherr *et al*., 2015) similarly offer conceptual parallels but typically require more complex mechanistic assumptions. Our approach aligns with these multiphase modeling strategies but is notably simpler, entirely data-driven, and implicitly captures biological adaptation without explicitly modeling regulatory or proteomic shifts.

Furthermore, dAMN uniquely predicts complete growth curves across unseen media, overcoming limitations of previous hybrid models. All the hybrid approaches mentioned in the introduction either lack generalization to new nutrient contexts, omit full dynamic modeling of biomass and metabolites, or rely on rich omics datasets not always available in experimental workflows. dAMN stands out in that it uses only initial metabolite concentrations, applies physics-inspired constraints, and can accurately extrapolate to forecast trajectories and to combinatorial media not seen during training.

## Author contribution

J.L.F. designed and supervised the study, acquired the fundings, and wrote the dAMN and dFBA codes. D.D. run the PINN code, created and maintained the GitHub implementation, formatted, tested and commented all codes. P.A. acquired the *E. coli* M28 experimental dataset. A.R. acquired the *P. putida* dataset under the supervision of E.A.G.. All authors contributed to writing, reading and editing the manuscript. All authors approved the manuscript.

## Acknowledgements

The authors would like to thank Alexandra Martin (INRAE, University of Paris Saclay) for her help in acquiring *E. coli* growth curves for different media and Anne Giralt (INRAE, University of Paris Saclay), Elouan Julien (Ecole Polytechnique) and Lucie-Garance Barot (ENS-Paris) for the development of PINN software used in the current study.

## Code and Data Availability

The python source codes associated to the paper the data and the notebook used to produce the results are available at https://github.com/brsynth/dAMN-main-release and DOI 10.5281/zenodo.17908125. All training data can be found in the repository in the folder ‘data’, trained models can be found in the folder ‘model’. The ‘data’ folder contains the file ‘Datasets.xls’ will all experimental datasets used in the study. Additional materials (text, figures and tables) are provided in the Supplementary Information file (SI).

