## Supplementary Information for "dAMN: a genome scale neural-mechanistic hybrid model to predict bacterial growth dynamics"

—

### Supplementary Information

Jean-Loup Faulon[1,*], Danilo Dursoniah[1], Paul Ahavi[1], Antoine Raynal[2], Enrique Asin-Garcia[2]
[1]University Paris Saclay, INRAE, AgroParisTech, MICALIS Institute, 78350 Jouy-en-Josas, France.
[2] Bioprocess Engineering Group, Wageningen University & Research, Wageningen, 6700 AA, The Netherlands

#### Table of content

1. dAMN model architecture and parameters 2
 1.1. Architectures 2
 Fig. S1. dAMN architectures. 2
 Fig. S2. dAMN results with Hill function for lag-phase. 3
 1.2. Hyperparameters 3
 Table S1. dAMN loss function hyperparameter search. 4
 Table S2. dAMN performances for different train-test splits. 4
 Fig. S3. dAMN results using prior loss without time decay. 5

2. *E. coli* M28 dataset 6
 2.1. Experimental methods 6
 2.2. dAMN results 7
 Fig. S4. Poor performing examples. 7
 Fig. S5. Predicting different lag phase times. 7

3. *P. putida* KT2440 dataset 8
 3.1. Experimental methods 8
 3.2. dAMN results 10
 Fig. S6. dAMN results with *P. putida* dataset. 10

4. *E. coli* Millard dataset 11
 4.1. Acetate overflow 11
 4.2. ODE model and measured concentration 11
 4.3. PINN architecture 12
 Table S3. Unknown kinetics parameters and search ranges for Millard ODE model. 13
 4.4. dAMN and PINN results 13
 Fig. S7. dAMN and PINN predictions for acetate overflow. 14
 Fig. S8. dAMN predictions for glucose-acetate consumption shift. 15

5. dFBA results with *E. coli* M28 dataset 16
 Fig. S9. dFBA predicted growth curves for the *E. coli* M28 dataset. 17

6. Supplementary references 17

### 1. dAMN model architecture and parameters

#### 1.1. Architectures

We developed two architectures for dAMN. These architectures differ only the way the lag-phase is being modeled. The first one (Fig. S1 panel (a)) is using an exponential function, and is described in the main manuscript (*cf*. Eqs. 1-12). The second is an alternative architecture where the lag phase is modeled using a Hill function. This alternative, already used with dFBA (Flassig *et al.*, 2016) is shown in Fig. S1 panel (b). We note that, with this alternative architecture, only one parameter $t_{lag}$ is learned as the stiffness parameter $n$ was set to 4 an in (Flassig *et al.*, 2016).

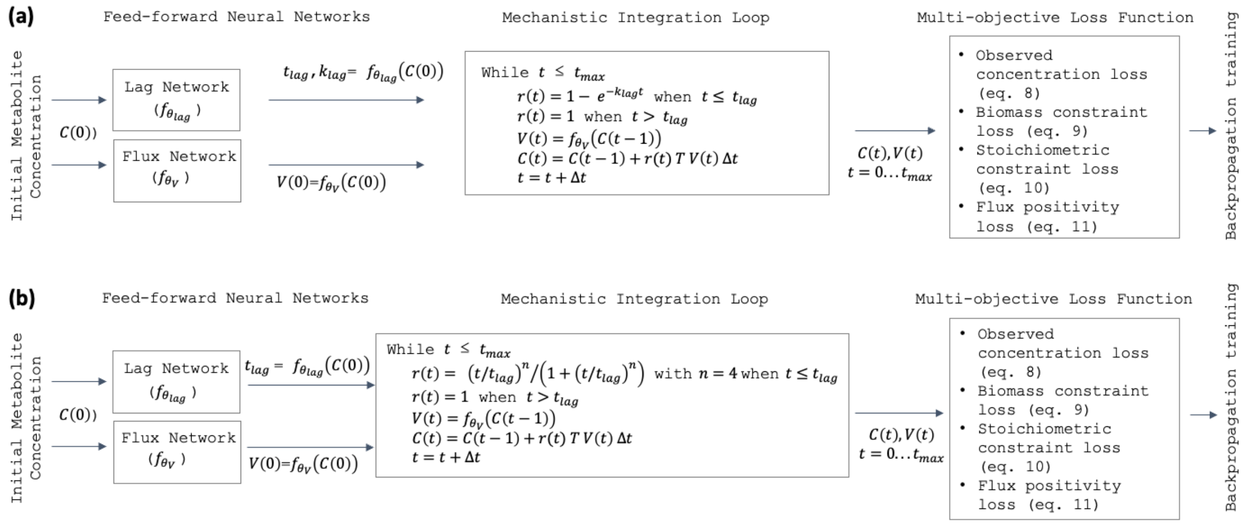


**Fig. S1. dAMN architectures.**

*(a) dAMN hybrid architecture is composed of two neural networks (to predict lag time parameters and fluxes) and followed by mechanistic equations iterated from t=0 to t=$t_{max}$. The lag network ($f_{\theta_{lag}}$), takes the initial concentration vector C(0) of medium metabolites as input, has a single hidden layer of 50 neurons with a ReLU activation function, and an output layer of size 2, predicting lag-phase parameters (lag time $t_{lag}$ and stiffness $k_{lag}$). A dropout rate of 0.2 is applied for regularization. To ensure output stability and positivity, both outputs are processed using a softplus activation function. The second neural network ($f_{\theta_V}$), the flux network, takes the concentration vector C(t) as input and predict fluxes for all reactions. It includes a hidden layer consisting of 500 neurons with ReLU activation and an output layer with a size equal to the number of reactions of the genome scale model (3682 for E. coli and 4034 for P. putida). Dropout regularization with a rate of 0.2 is employed, and linear activation is used in the output layer to predict continuous flux values. (b) This architecture is composed of two neural networks (to predict lag time parameters and fluxes) and followed by the same mechanistic as in panel (a). The lag network ($f_{\theta_{lag}}$), takes the initial concentration vector C(0) of medium metabolites as input, has a single hidden layer of 50 neurons with a ReLU activation function, and an output layer of size 1, predicting lag time $t_{lag}$.*

The results obtained with the alternative architecture (Hill function) for the M28 *E. coli* data set with the forecasting training are presented in Fig. S2. These are similar to those obtained with the exponential function (*cf*. Figure 1 in main manuscript).

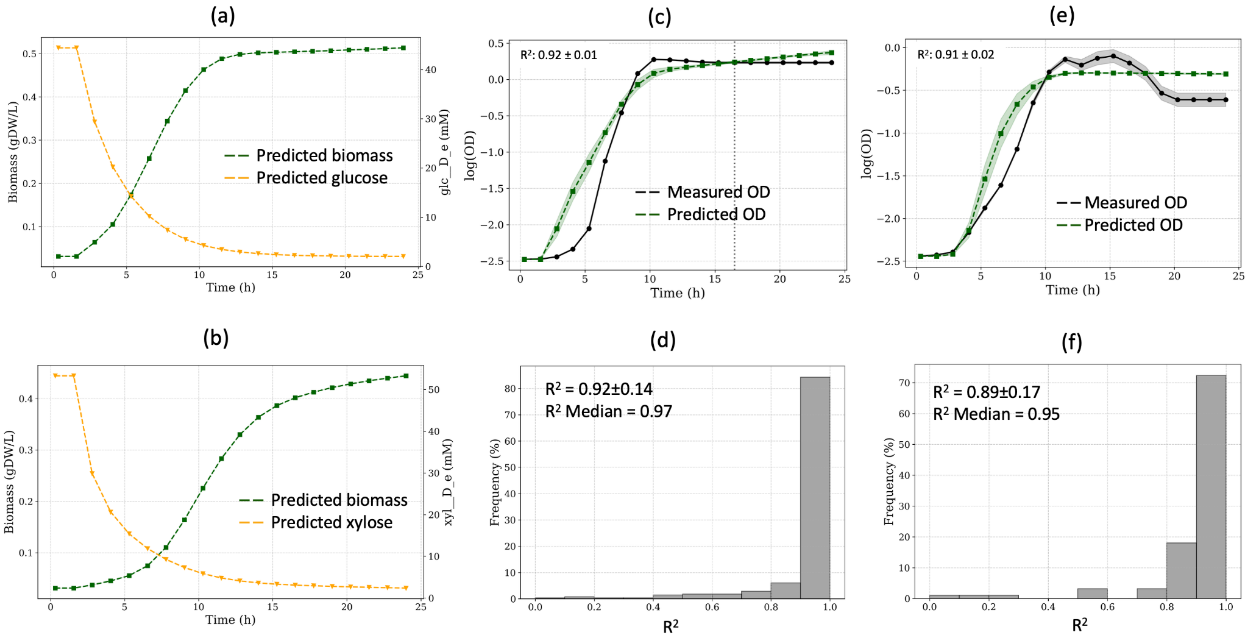


**Fig. S2. dAMN results with Hill function for lag-phase.**

*(a) Predicted concentrations for biomass and D-glucose when the strain is grown on M9 with D-glucose only. (b) Predicted concentrations for biomass and xylose when the strain is grown on M9 with xylose only. (c) Example of a predicted growth curve compared to measured growth curve using the forecast training set (dashed line indicates when forecast begins). dAMN is trained on the first two-thirds of the datapoints, with predictions made for the final third. (d) Distribution of $R^2$ values between measured and predicted growth curves for the forecast training set. (e) Example of a predicted growth curve compared to measured growth curve using the media training set. dAMN is trained on the first two-thirds of all media, and predictions are made for media not used during training. The displayed curve represents prediction for a medium not present in the training set. (f) Distribution of $R^2$ values between measured and predicted growth curves for the media training set. For all plots, the shaded area represents standard deviation obtained for 3 repeats.*

#### 1.2. Hyperparameters

The dAMN loss function is composed of 4 terms (*cf*. Eqs.7-11 in main manuscript). Each of these four terms are weighted by an exponential decay function which evolves over time (Eq. 12 in main manuscript). The parameters of the decay functions consist of a weight $\lambda_i$ and a decay $k_i$. To search for the parameters maximizing dAMN predictability, we conducted a parameter grid search using the M28 *E. coli* dataset with the forecasting training set. Results are presented in Table S1.

Using the best parameter set of Table S1, we trained dAMN using different validation folds and different train-test splits. Results are presented in Table S2.

**Table S1. dAMN loss function hyperparameter search.**

*The parameters correspond to those in Eq. 12. To limit the search space, $\lambda_1$, $\lambda_2$ and $\lambda_4$ were fixed at 1, and $k_3$, the decay parameter for the stoichiometric constraint, was set to 0, ensuring that the stoichiometric constraint remained constant over time. Variances for $R^2$-mean are not single growth curve variances but are variances obtained for the 280 growth curves.*

| $\lambda_1$ | $\lambda_2$ | $\lambda_3$ | $\lambda_4$ | $k_1$ | $k_2$ | $k_3$ | $k_4$ | R2-mean | R2-median |
|---|---|---|---|---|---|---|---|---|---|
| 1 | 1 | 0.1 | 1 | 0 | 0 | 0 | 1 | 0.84±0.2 | 0.91 |
| 1 | 1 | 0.001 | 1 | 0.5 | 1 | 0 | 0.5 | 0.84±0.2 | 0.91 |
| 1 | 1 | 0.001 | 1 | 0.25 | 0.5 | 0 | 0.25 | 0.83±0.22 | 0.91 |
| 1 | 1 | 0.01 | 1 | 0.5 | 1 | 0 | 0.75 | 0.82±0.2 | 0.90 |
| 1 | 1 | 0.1 | 1 | 0 | 1 | 0 | 0.5 | 0.86±0.14 | 0.89 |
| 1 | 1 | 0.001 | 1 | 0.75 | 1 | 0 | 0.5 | 0.74±0.3 | 0.88 |
| 1 | 1 | 0.001 | 1 | 0.75 | 1 | 0 | 1 | 0.67±0.3 | 0.83 |
| 1 | 1 | 0.01 | 1 | 0.5 | 0.5 | 0 | 0.25 | 0.74±0.2 | 0.83 |
| 1 | 1 | 1 | 1 | 0.25 | 0.5 | 0 | 0.5 | 0.72±0.26 | 0.82 |
| 1 | 1 | 0.001 | 1 | 1 | 1 | 0 | 0.75 | 0.67±0.3 | 0.79 |
| 1 | 1 | 1 | 1 | 0.25 | 0 | 0 | 1 | 0.65±0.31 | 0.79 |
| 1 | 1 | 0.001 | 1 | 0.75 | 0 | 0 | 0.25 | 0.68±0.29 | 0.79 |
| 1 | 1 | 0.001 | 1 | 1 | 0.75 | 0 | 0.25 | 0.65±0.2 | 0.77 |
| 1 | 1 | 0.1 | 1 | 0.5 | 0.25 | 0 | 0.25 | 0.65±0.3 | 0.76 |
| 1 | 1 | 0.01 | 1 | 1 | 0 | 0 | 0 | 0.62±0.3 | 0.76 |
| 1 | 1 | 1 | 1 | 0.5 | 0.75 | 0 | 1 | 0.64±0.27 | 0.76 |
| 1 | 1 | 1 | 1 | 0.75 | 1 | 0 | 0.5 | 0.48±0.28 | 0.54 |
| 1 | 1 | 1 | 1 | 0.5 | 0 | 0 | 0.75 | 0.33±0.31 | 0.40 |
| 1 | 1 | 0.1 | 1 | 1 | 0 | 0 | 0.25 | 0.24±0.26 | 0.14 |
| 1 | 1 | 1 | 1 | 0.75 | 0 | 0 | 0 | 0.23±0.25 | 0.13 |

**Table S2. dAMN performances for different train-test splits.**

*dAMN was run with the M28 E. coli dataset with medium training sets of various size and train-test splits. The decay function parameters for the loss function are those of the first row of Table S1.*

| x-fold | Training size | Test size | R2 mean | R2 median | R2 mean | R2 median |
|---|---|---|---|---|---|---|
| 3 | 187 | 93 | 0.90 | 0.96 | 0.90±0.01 | 0.95±0.01 |
| 3 | 187 | 93 | 0.90 | 0.93 | | |

| | | | | | | |
|---|---|---|---|---|---|---|
| 3 | 187 | 93 | 0.89 | 0.95 | | |
| 2 | 140 | 140 | 0.87 | 0.94 | 0.87±0.01 | 0.93±0.01 |
| 2 | 140 | 140 | 0.88 | 0.93 | | |
| 2 | 140 | 140 | 0.87 | 0.93 | | |
| 1.5 | 93 | 187 | 0.84 | 0.92 | 0.85±0.03 | 0.92±0.02 |
| 1.5 | 93 | 187 | 0.88 | 0.94 | | |
| 1.5 | 93 | 187 | 0.82 | 0.91 | | |

We also tested the ability of dAMN to predict growth curves without a time decay in the loss function. To that end, we set all $\lambda_i$ values to 1 and all $k_i$ values to 0. To obtain an equal contribution of each of the four losses we used the prior loss formulation where loss at epoch ($n$) is divided by the loss at the previous epoch ($n$-1) i.e., $\text{loss}_i(n) = \text{loss}_i(n)/\text{loss}_i(n-1)$ The results, presented in Fig. S3 panel (b), demonstrates that biomass depletion cannot be modeled with the prior loss.

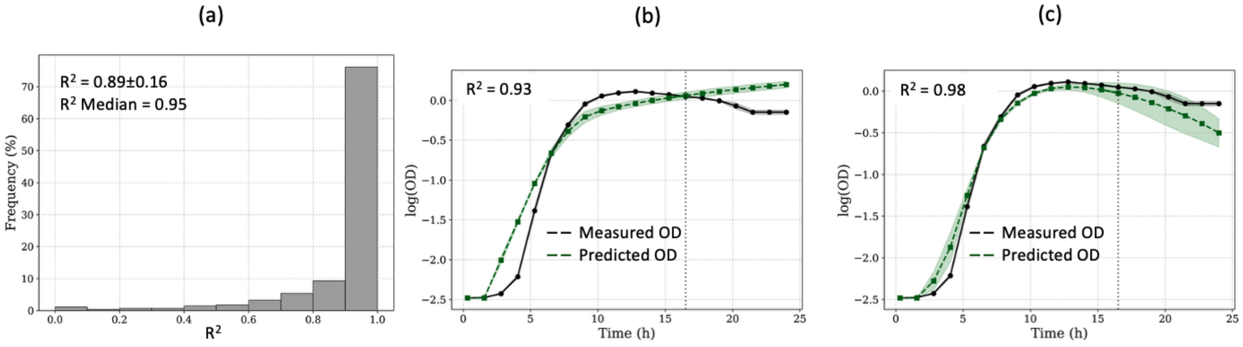


**Fig. S3. dAMN results using prior loss without time decay.**

*(a) Distribution of $R^2$ values between measured and predicted growth curves for the M28 forecast training set with a prior loss. (b) Example (medium number 168) of a predicted growth curve compared to measured growth curve using prior loss (dashed line indicates when forecast begins). dAMN is trained on the first two-thirds of the datapoints, with predictions made for the final third. (b) Same example (medium number 168) of a predicted growth curve compared to measured growth curve using exponential decay loss functions with parameters in the first row of Table S1. For all plots, the shaded area represents standard deviation obtained for 3 repeats.*

### 2. *E. coli* M28 dataset

#### 2.1. Experimental methods

***Bacterial strain and culture media.*** The strain used to construct the *E. coli* M28 dataset was *Escherichia coli* K-12 MG1655 (accession number NC_000913.3). The full procedure for building this dataset has been detailed previously in (Ahavi *et al.*, 2024).

In summary, the aim was to assemble a collection of media capable of generating a broad diversity of growth curves for *E. coli* K-12 MG1655. To this effect, an M9-based medium was supplemented with selected combinations of three carbon sources (D-glucose, D-xylose, or succinate), purine nucleobases, pyrimidine nucleobases, and the 20 proteinogenic amino acids grouped into nine biosynthetic classes (Gly/Ser/Thr; Asn/Asp/Ala; Glu/Gln; Arg/Pro; Val/Leu/Ile; Lys; Cys/Met; His; Phe/Tyr/Trp). These groups were ranked according to their estimated energetic contribution to the cell (Zampieri *et al.*, 2019). By varying both the number of amino acid groups added and the rank of each group, a wide range of growth phenotypes could be obtained. The three carbon sources differ in their energetic yield for *E. coli* and therefore further expanded the diversity of achievable growth behaviors. The composition of the M9-base medium and the nutrient concentrations are provided in (Ahavi *et al.*, 2024).

***Bacterial growth monitoring***. A detailed protocol has been described in (Ahavi *et al.*, 2024). Briefly, *E. coli* K-12 MG1655 was recovered from a glycerol stock onto an M9 agar plate supplemented with 44.4 mM (0.8%) D-(+)-glucose. The plate was incubated overnight at 37 °C. A single colony was grown in 10 mL of the corresponding liquid medium until the optical density at 600 nm reached ~0.5. Two microliters of the culture were used to inoculate 198 µL of each M28 medium in flat-bottom 96-well plates. Each medium was prepared in triplicate. The plates were then incubated at 37 °C in a microplate reader (BioTek Synergy HTX, Agilent Technologies) with continuous orbital shaking (807 cpm, 1 mm). Turbidity (optical density at 600 nm) was measured and recorded every 20 minutes for assays lasting between 16 and 24 hours.

All data (media metabolite concentration and OD measurements over time) are provided in Datasets.xls supplementary file available at DOI 10.5281/zenodo.17908125 in the 'data' folder.

#### 2.2. dAMN results

The results obtained with dAMN for the *E. coli* M28 dataset are presented in Figure 1 of the main manuscript. From these results we extracted in Fig. S4 cases with poor predictions, and in Fig. S5 cases with different lag times.

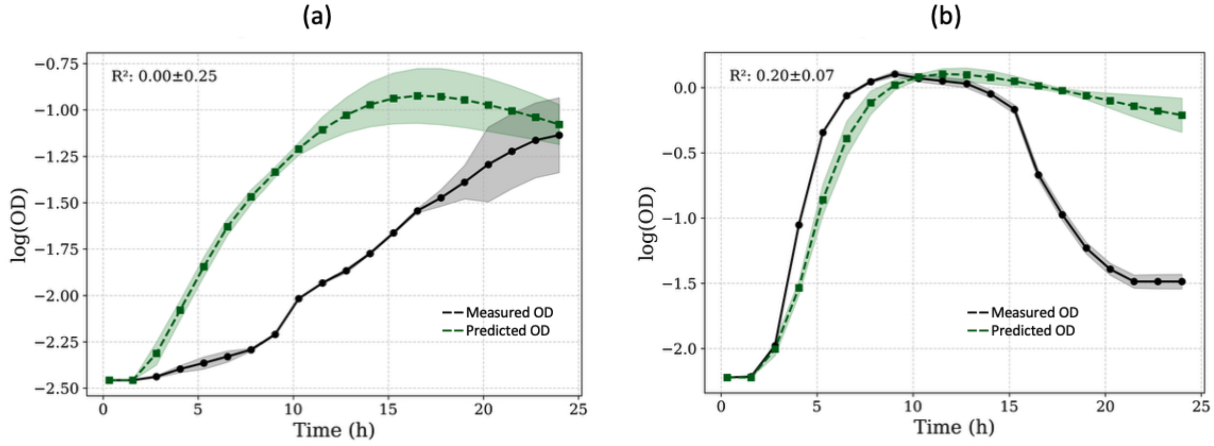


**Fig. S4. Poor performing examples.**

*dAMN was trained on the medium dataset used in Figure 2 (main manuscript). (a) An example with no obvious change between lag and exponential phases (medium number 280). (b) An example with sharp growth depletion (medium number 201). For all plots, the shaded area represents standard deviation obtained for 3 repeats.*

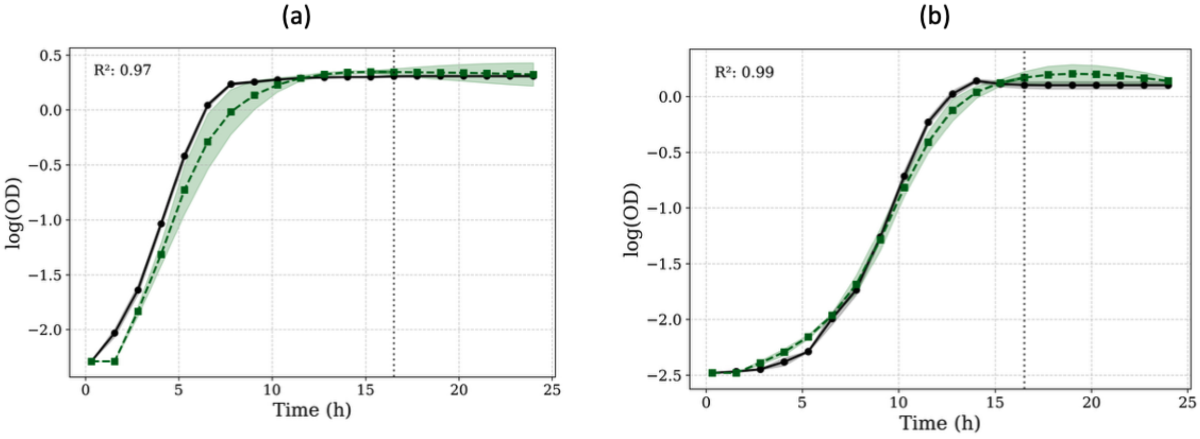


**Fig. S5. Predicting different lag phase times.**

*(a) An example with no lag phase medium number 72). (b) An example where lag phase extends up to > 5 hours (medium 84). For all plots, the shaded area represents standard deviation obtained for 3 repeats.*

### 3. *P. putida* KT2440 dataset

#### 3.1. Experimental methods

***Bacterial strain and culture media***. The strain used to construct the *P. putida* dataset was *Pseudomonas putida* KT2440 (accession number: NC_002947.4). All culture media were based on *P. putida* M9 minimal medium. The M9 medium was prepared in 940 mL sterile deionized $H_2O$ and supplemented with the following sterile stock solutions: 20 mL of 50x phosphate buffer (pH7) consisting of 388 g/L $K_2HPO_4$ (Merck/Sigma-Aldrich, CAS 7758-11-4) and 163 g/L $NaH_2PO_4$ (Merck/Sigma-Aldrich, CAS 10049-21-5); 20 mL of 50x ammonium sulfate solution (pH7) containing 200 g/L $(NH_4)_2SO_4$ (Merck/Sigma-Aldrich, CAS 7783-20-2); 20 mL of 50x salts solution (pH7) containing 1 g/L EDTA (Merck/Sigma-Aldrich, CAS 60-00-4), 10 g/L $MgCl_2 \cdot 6H_2O$ (Merck/Sigma-Aldrich, CAS 7791-18-6), 0.2 g/L $ZnSO_4 \cdot 7H_2O$ (Merck/Sigma-Aldrich, CAS 7446-20-0), 0.1 g/L $CaCl_2 \cdot 2H_2O$ (Merck/Sigma-Aldrich, CAS 10035-04-8), 0.5 g/L $FeSO_4 \cdot 7H_2O$ (Merck/Sigma-Aldrich, CAS 7782-63-0), 0.02 g/L $Na_2MoO_4 \cdot 5H_2O$ (Merck/Sigma-Aldrich, CAS 10102-40-6), 0.02 g/L $CuSO_4 \cdot 5H_2O$ (Merck/Sigma-Aldrich, CAS 7758-99-8), 0.04 g/L $CoCl_2 \cdot 6H_2O$ (Thermo Fisher Scientific, CAS 10026-22-9), and 0.1 g/L $MnCl_2 \cdot 4H_2O$ (Merck/Sigma-Aldrich, CAS 13446-34-9). After addition of the stock solutions, the final volume of the M9 minimal medium was adjusted to 1 L.

As with *E. coli (*section 2.1), this base medium was supplemented with different combinations of nutrients selected from 4 sugars or organic acids, 20 proteinogenic amino acids, and 5 nucleobases. All stock solutions were filter-sterilized using 0.2 µm membranes. Carbon sources were added to the following final concentrations: 3.96 g/L D-(+)-glucose (Thermo Fisher Scientific, CAS 14431-43-7), 3.6 g/L D-(-)-fructose (Merck/Sigma-Aldrich, CAS 57-48-7), 0.59 g/L acetate (Merck/Sigma-Aldrich, CAS 127-09-3), and 1.16 g/L succinate (Merck/Sigma-Aldrich, CAS 6106-21-4). Additional combinations included 1.89 g/L citrate (Merck/Sigma-Aldrich, CAS 77-92-9), 1.34 g/L malate (Merck/Sigma-Aldrich, CAS 6915-15-7), or 0.87 g/L pyruvate (Merck/Sigma-Aldrich, CAS 617-35-6).

Amino acids and nucleobases were added to the following final concentrations: 0.89 g/L L-alanine (Merck/Sigma-Aldrich, CAS 56-41-7), 1.74 g/L L-arginine (Merck/Sigma-Aldrich, CAS 74-79-3), 1.32 g/L L-asparagine (Merck/Sigma-Aldrich, CAS 70-47-3), 1.33 g/L L-aspartic acid (Merck/Sigma-Aldrich, CAS 56-84-8), 0.24 g/L L-cysteine (Merck/Sigma-Aldrich, CAS 52-90-4),

1.47 g/L L-glutamic acid (Merck/Sigma-Aldrich, CAS 56-86-0 ), 1.46 g/L L-glutamine (Merck/Sigma-Aldrich, CAS 56-85-9), 0.75 g/L glycine (Merck/Sigma-Aldrich, CAS 56-40-6), 0.77 g/L L-histidine (Merck/Sigma-Aldrich, CAS 5934-29-2), 1.31 g/L L-isoleucine (Merck/Sigma-Aldrich, CAS 73-32-5), 1.31 g/L L-leucine (Merck/Sigma-Aldrich, CAS 61-90-5), 0.73 g/L L-lysine (Merck/Sigma-Aldrich, CAS 657-27-2), 0.75 g/L L-methionine (Merck/Sigma-Aldrich, CAS 63-68-3), 1.65 g/L L-phenylalanine (Merck/Sigma-Aldrich, CAS 63-91-2), 1.15 g/L L-proline (Merck/Sigma-Aldrich, CAS 147-85-3), 1.05 g/L L-serine (Merck/Sigma-Aldrich, CAS 56-45-1), 1.19 g/L L-threonine (Merck/Sigma-Aldrich, CAS 72-19-5), 1.02 g/L L-tryptophan (Merck/Sigma-Aldrich, CAS 73-22-3), 0.91 g/L L-tyrosine (Merck/Sigma-Aldrich, CAS 60-18-4), 1.17 g/L L-valine (Merck/Sigma-Aldrich, CAS 72-18-4), 0.27 g/L adenine (Merck/Sigma-Aldrich, CAS 321-30-2 ), 0.22 g/L cytosine (Thermo Fisher Scientific, CAS 71-30-7 ), 0.22 g/L uracil (Merck/Sigma-Aldrich, CAS 66-22-8), 0.30 g/L guanine (Thermo Fisher Scientific, CAS 73-40-5), and 0.48 g/L thymidine (Thermo Fisher Scientific, CAS 50-89-5).

In summary, the aim was to assemble a collection of media capable of generating a broad diversity of growth curves for *P. putida* KT2440. Nutrient combinations were designed to impose specific biosynthetic demands, including nitrogen metabolism (Glu/Gln), urea cycle and alkaline amino acids (Arg/Lys/Pro), sulfur and methyl cycle load (Cys/Met), aromatic biosynthetic burden (Phe/Tyr/Trp), branch-chained amino acids (Ile/Leu/Val), and pentose phosphate-linked histidine synthesis. Combination of small amino acids (Gly/Ser/Thr; Ala/Asp/Asn) targeted transamination and one-carbon metabolism.

***Bacterial growth monitoring***. A glycerol stock of *P. putida* KT2440 was streaked onto LB agar and incubated overnight at 30 °C. A single colony was used to inoculate a preculture of 10 mL of LB medium in a 50 mL Falcon tube, followed by overnight incubation at 30 °C with shaking at 250 rpm (New Brunswick™ Innova®42/42R). 20 µL of this culture was then transferred to M9 minimal medium supplemented with 20 mM D-(+)-glucose and incubated overnight under the same conditions. For inoculum preparation, three 700 µL aliquots of the culture were washed twice in unsupplemented M9 without centrifugation at 7,000 rcf for 1 min, discarding the supernatant each time, and resuspending to a final volume of 400 µL. Optical density at 600 nm was measured using a DR6000 UV-VIS spectrophotometer (Hach Lange), and the suspension was diluted to obtain an $OD_{600}$ of 0.1 when 5 µL were added to a final volume of 200 µL. For the growth assays, 195 µL of each nutrient combination was dispensed into wells of a clear, flat-bottom 96-well plate and inoculated with the adjusted *P. putida* preculture to achieve a starting

$OD_{600}$ of approximately 0.1. All conditions were tested in triplicate. Plates were incubated at 30 °C in a BioTek HTX Synergy plate reader (Agilent Technologies) with continuous orbital shaking. Optical density at 600 nm was recorded every 15 min for 24-48 h to monitor bacterial growth.

All data (media metabolite concentration and OD measurements over time) are provided in Datasets.xls supplementary file available at DOI 10.5281/zenodo.17908125 in the 'data' folder.

##### 3.2. dAMN results

We created two training sets with *P. putida* following the same procedure as for the *E. coli* M28 dataset. In the first set, the forecast set, dAMN is trained using two-thirds of the growth time points from all available data, and predictions are made on the remaining one-third of the time points. In the second set, the media set, dAMN is trained on two-thirds of the available media using all time points, and predictions are made for media conditions not included in the training set. Fig. S6 below shows that the results obtained are similar to those obtained with *E. coli*, despite the fact that the *P. putida* dataset is much smaller (81 media instead of 280).

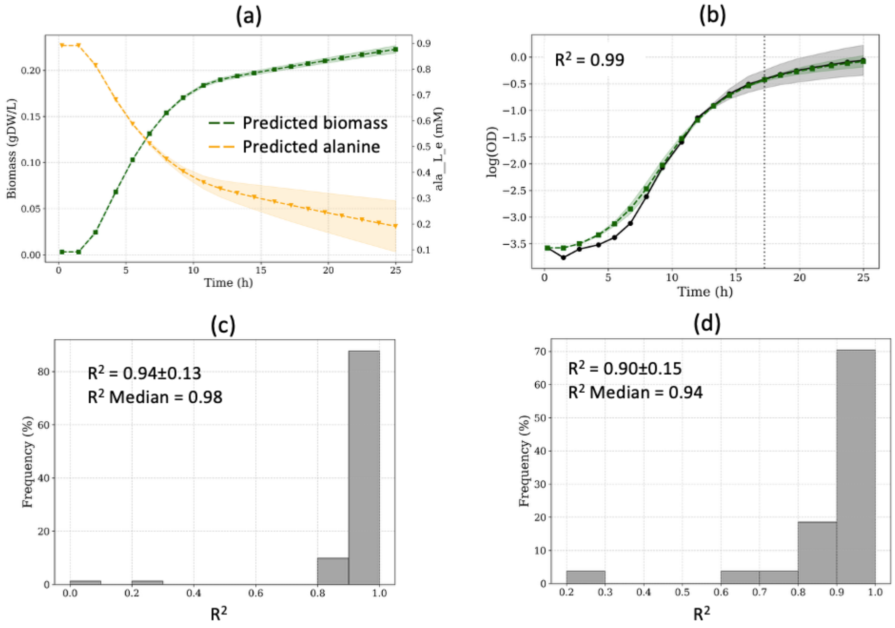


**Fig. S6. dAMN results with *P. putida* dataset.**

*(a) Predicted concentrations for biomass and alanine when the strain is grown on M9 with only alanine as carbon source (medium number 81). (b) Example of prediction in forecasting mode (medium number 26, dashed line indicates when forecast begins). dAMN is trained on the first two-thirds of the datapoints, with predictions made for the final third. (c) Distribution of $R^2$ values between measured and predicted growth curves for the forecast training set. (d) Distribution of $R^2$ values between measured and predicted growth curves for the media training set. For all plots, the shaded area represents standard deviation obtained for 3 repeats.*

### 4. *E. coli* Millard dataset

#### 4.1. Acetate overflow

Acetate overflow refers to the phenomenon in which *E. coli* secretes acetate during growth on excess glucose, even under fully aerobic conditions. It arises when the cellular capacity to generate acetyl-CoA via glycolysis exceeds the capacity of the TCA cycle and biosynthetic pathways to consume it, causing carbon to be diverted through the Pta–AckA pathway to acetate. This imbalance is further shaped by thermodynamic control of the acetyl-CoA/acetate node and by global regulatory responses that down-modulate glycolysis and the TCA cycle as extracellular acetate accumulates. In the dataset extracted from (Millard *et al.*, 2021) and used in the current study, *E. coli* K-12 MG1655 was grown in M9 medium complemented with 15 mM glucose. Sodium acetate (prepared in a 1.6 M solution at pH 7.0) was added up to 1mM concentration. All data (media metabolite initial concentration and metabolite concentrations over time) are provided in Datasets.xls supplementary file available at DOI 10.5281/zenodo.17908125 in the 'data' folder.

#### 4.2. ODE model and measured concentration

In (Millard *et al.*, 2021) the dynamic model of *E. coli* glucose–acetate metabolism is formulated as a system of six ordinary differential equations (ODEs) describing extracellular glucose (GLC), extracellular acetate ($ACE_{env}$), biomass (X), and intracellular metabolites acetyl-CoA (AcCoA), acetyl-phosphate (AcP), and intracellular acetate ($ACE_{cell}$). Each state variable evolves according to mass-balance equations driven by the rates of glycolysis, the TCA cycle, the Pta–AckA pathway, and acetate exchange. In the general model, glycolysis and the TCA cycle follow irreversible Michaelis–Menten kinetics with optional non-competitive inhibition by extracellular acetate; the Pta and AckA reactions are described by reversible Michaelis–Menten laws; and acetate exchange follows reversible saturation kinetics. Growth is computed directly from the TCA flux via a constant biomass yield. The complete equation set is:

$$
\begin{aligned}
&dGLC/dt = v_{glycolysis}\ X\ V_{cell}/V_{env} + v_{feed} - D\ GLC \\
&dACE_{env}/dt = v_{acetate\ exchange}\ X\ V_{cell}/V_{env} - D\ ACE_{env} \\
&dX/dt = X\ v_{growth} - DX \\
&dACCOA/dt = 1.4\ v_{glycolysis} - v_{TCA\ cycle} - v_{Pta} \\
&dACP/dt = v_{Pta} - v_{AckA} \\
&dACE_{cell}/dt = v_{AckA} - v_{acetate\ exchange}
\end{aligned}
$$

Each flux $v_i$ depends on a set of kinetic parameters (e.g., $V_{max}$, $K_m$, inhibition constants $K_i$, and equilibrium constants $K_{eq}$), whose values are taken from literature or estimated from $^{13}$C-labelling time-courses. The model outputs (observables) are extracellular concentrations of glucose, acetate, and biomass.

In the present study, we use exclusively the parameterization and experimental dataset corresponding to the 1 mM acetate condition, i.e., the lowest acetate background. This regime avoids confounding effects of acetate-induced inhibition on glycolysis and the TCA cycle and reflects the baseline overflow phenotype under minimal perturbation. All simulations, fits, and comparisons therefore rely on the 1 mM condition as the representative dynamical system.

##### 4.3. PINN architecture

We used a conventional physics-informed neural network (PINN) following the formulation of (Raissi *et al.*, 2019) used in many ODE- and PDE-based studies. In our implementation, the PINN maps time *t* to the six Millard state variables (GLC, $ACE_{env}$, X, ACCOA, ACP, $ACE_{cell}$). The network is a fully-connected multi-linear perceptron (MLP):

- input: time *t*;
- hidden layers: 7 layers × 20 neurons, Softplus activation;
- output: 6 metabolic variables;
- weight initialization: Xavier uniform.

Training minimizes a weighted sum of three components (total loss is: $L = L_{data} + L_{ode} + L_{aux}$) :

- Data-fit loss $L_{data}$ on observed variables (GLC, $ACE_{env}$, X): $L_{data} = \Sigma_j \,||\, \hat{y}_j(t) - y_j(t) \,||^2$ , with $\hat{y}_j(t)$ and $y_j(t)$ the PINN predicted trajectory and the experimental measurements for the observable $j$, respectively.
- Physics (ODE-residual) loss $L_{ode}$ enforcing the Millard model: $L_{ode} = \Sigma_i \,||\, \frac{d\hat{y}_i}{dt} - f_i(\hat{y}, \hat{\theta}) \,||^2$ using automatic differentiation. $i$ denotes any state variable (observable or latent), hence, $\hat{y}_i$ and $y_i$ denotes the PINN predicted trajectory and the experimental measurements for the observable $i$, respectively, while $f_i$ is the Millard ODE right-hand side for this state variable integrating the learned kinetic parameters, $\hat{\theta}$.

- Auxiliary loss $L_{aux}$ constraining initial and final states: $L_{aux} = \Sigma_i [(\hat{y}_i(t_0) - y_i^0)^2 + (\hat{y}_i(t_f) - y_i^f)^2]$, where $y_i^0$ and $y_i^f$ are the initial and final values of a variable $i$, at initial time $t_0$ and final time $t_f$, respectively.

In addition, we used a simple prior-loss reweighting heuristic to improve numerical stability when combining heterogeneous loss terms (data, residuals, and auxiliary constraints). For each component $(L_i)$, a weight is computed from its current and earlier epochs values - respectively $n$ and $n$-1: $L_i^{(n)} = L_i^{(n)} / L_i^{(n-1)}$

As in (Yazdani *et al.*, 2020) unknown kinetic parameters ($\hat{\theta}$) were learned jointly with the network. These kinetic parameters are listed in Table S3.

**Table S3. Unknown kinetics parameters and search ranges for Millard ODE model.**

This *Table is adapted from (Millard et al., 2021). The exact parameter values (Value column) were computed in Millard et al. using a Monte Carlo approach.*

| Reaction | Parameter | Range | Value | Unit |
|---|---|---|---|---|
| AckA | $V_{max}$ | $2.8 \times 10^5 – 5.5 \times 10^5$ | 336940.02 | $mmol \cdot L^{-1} \cdot h^{-1}$ |
| Pta | $V_{max}$ | $4.9 \times 10^4 – 9.9 \times 10^6$ | 9565521.76 | $mmol \cdot L^{-1} \cdot h^{-1}$ |
| Glycolysis | $V_{max}$ | $5.3 \times 10^3 – 5.9 \times 10^3$ | 5557.64 | $mmol \cdot L^{-1} \cdot h^{-1}$ |
| | $Ki_{ACE}$ | 30.9 – 46.9 | 36.6776 | mM |
| TCA cycle | $Km_{COA}$ | 8.4 – 615.4 | 24.76 | mM |
| | $V_{max}$ | $2.4 \times 10^5 – 1.7 \times 10^6$ | 740859.87 | $mmol \cdot L^{-1} \cdot h^{-1}$ |
| | $Ki_{ACE}$ | 1.8 – 3.4 | 2.33 | mM |
| Growth | $Y$ | $9 \times 10^5 – 1.1 \times 10^{-4}$ | 9.98e-05 | $gDW.mmol^{-1}$ |
| Acetate exchange | $V_{max}$ | $8 \times 10^4 – 1.5 \times 10^6$ | 480034.55 | $mmol \cdot L^{-1} \cdot h^{-1}$ |
| | $Km_{ACE}$ | 1.5 – 99.8 | 33.15 | mM |

The raw trainable parameters were mapped to bounded ranges using a tanh-based transformation to ensure physiological plausibility. For training we used the Adam optimization algorithm with a cyclic learning-rate scheduler, as customary with PINN.

##### 4.4. dAMN and PINN results

The Millard data set was used with dAMM and PINN. In both cases training was performed on (i) acetate extracellular and biomass concentrations and (ii) glucose and biomass concentrations. Predictions were respectively made for glucose (i) and extracellular acetate (ii) concentrations. Results are shown in Fig. S7. We note that PINN struggles to predict glucose or acetate concentrations (panels (c) and (d)) when kinetics parameters are inferred by the PINN from the ranges provided in Table S3, but PINN predictions are similar to those obtained with dAMN when extract parameters values are provided (panels (e) and (f))

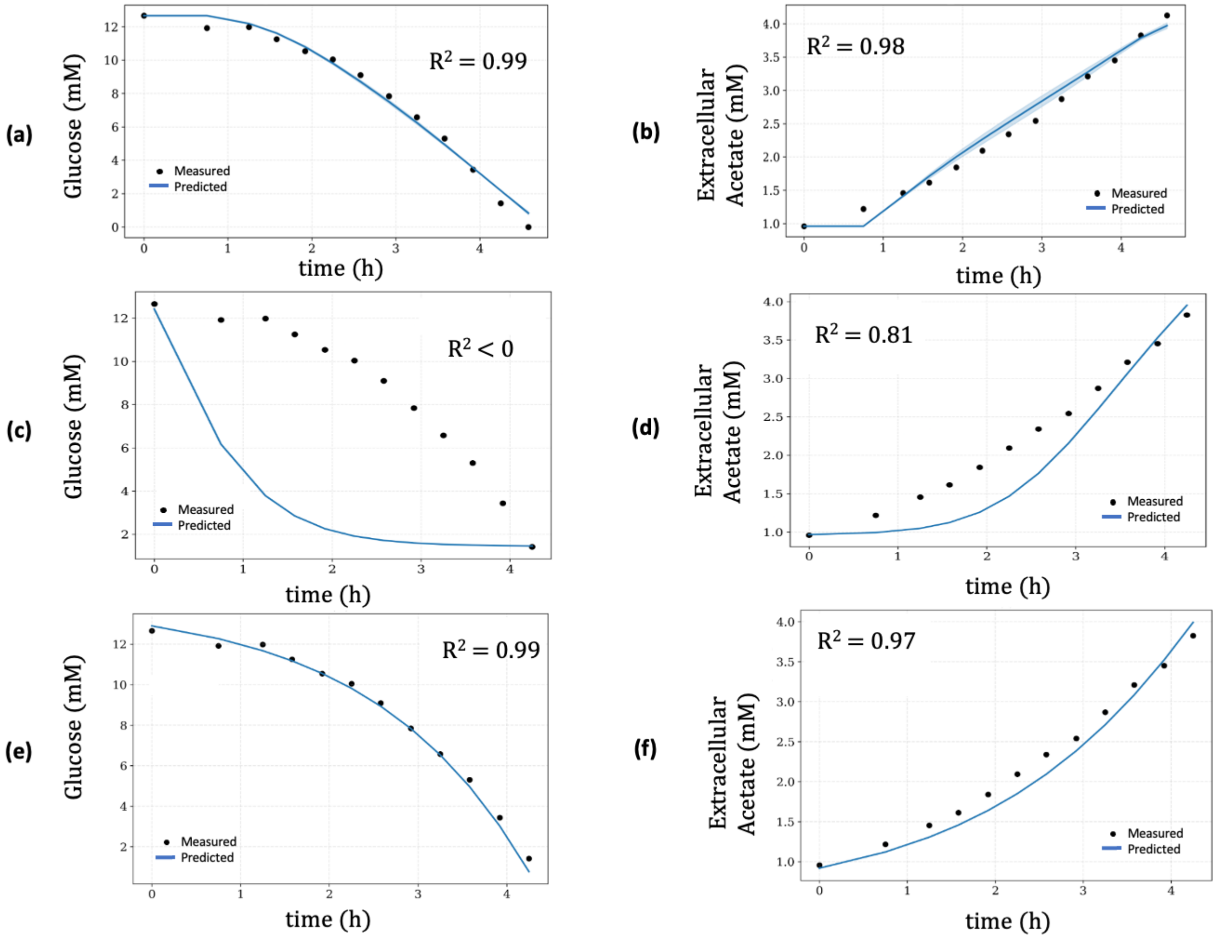


**Fig. S7. dAMN and PINN predictions for acetate overflow.**

*(a) Glucose concentration prediction when dAMN is trained on extracellular acetate and biomass concentrations. (b) Extracellular acetate concentration prediction when dAMN is trained on glucose and biomass concentrations. (c) Glucose concentration prediction for PINN trained on extracellular acetate and biomass concentrations with kinetics parameters searched in the range provided in Table S3 (d) Extracellular acetate concentration prediction for PINN trained on glucose and biomass concentrations with kinetics parameters searched in the range provided in Table S3. (e) Glucose concentration prediction for PINN trained on extracellular acetate and biomass concentrations with exact kinetics parameters values provided in Table S3. (f) Extracellular acetate concentration prediction for PINN trained on glucose and*

*biomass concentrations with exact kinetics parameters values provided in Table S3. For all plots, the shaded area represents standard deviation obtained for 3 repeats.*

To evaluate whether dAMN can capture multi-substrate dynamics, we extended the simulation beyond the experimental time window. Precisely, we trained dAMN on glucose and biomass concentrations up to 4.5 hrs and predicted the concentrations of glucose, acetate and biomass up to 9 hrs. As shown in Fig. S8, the model predicted acetate overflow (as in Fig. S7) but also subsequent acetate consumption and continued biomass increase, consistent with classical diauxic behavior even though these phases are not observed in the experimental data.

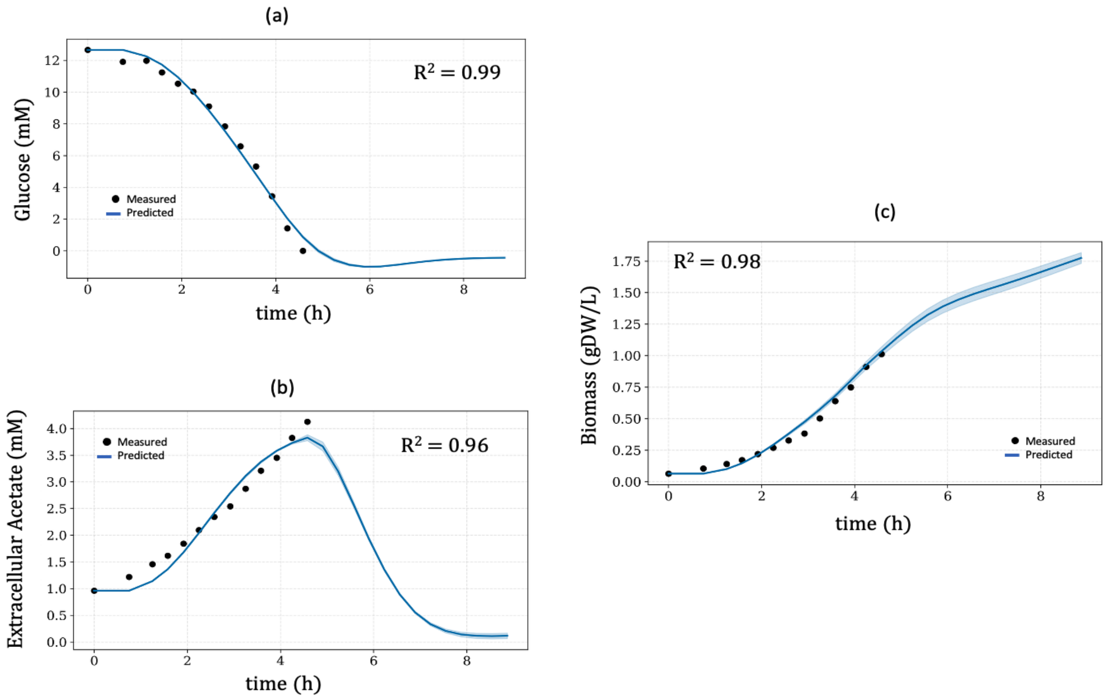


**Fig. S8. dAMN predictions for glucose-acetate consumption shift.**

*In all cases experimental data are provided up to 4.25 hrs and predictions are made up to 9 hrs. dAMN is trained on glucose and biomass measured data, acetate measured concentrations are not used in training. (a) Glucose consumption (b) Acetate overflow and consumption. (c) Biomass concentration. For all plots, the shaded area represents standard deviation obtained for 3 repeats.*

#### 5. dFBA results with *E. coli* M28 dataset

To benchmark mechanistic model performance, we simulated all 280 media conditions of the M28 *E. coli* dataset using classical dynamic flux balance analysis (dFBA) implemented in COBRApy (Ebrahim *et al.*, 2013) (*cf*. https://cobrapy.readthedocs.io/en/latest/dfba.html). For each medium, the genome-scale model iML1515 was loaded, and exchange reactions associated with available extracellular metabolites were dynamically constrained at each time point. Specifically, for every exchange reaction $i$ the maximum allowable uptake flux was set based on the metabolite concentration measured at time $t$: $v_{max,i} = \alpha_i \, C_i(t)/(\Delta t \, V)$ where $C_i(t)$ is the extracellular concentration, $\Delta t$ the integration step, $V$ the culture volume, and $\alpha_i$ a per-metabolite uptake scaling coefficient. This formulation constrains nutrient consumption using a concentration-dependent bound while preserving the native reaction directionality in the metabolic model. At each step, a steady-state FBA problem was solved to obtain the instantaneous growth rate $\mu(t)$, after which extracellular concentrations and biomass were updated using mass-balance equations (*cf*. Eq. 5 in main manuscript). Initial biomass values were taken directly from the experimental OD measurements for each condition.

Two uptake-scaling coefficients $\alpha_i$ were used: one shared among sugars (glucose, xylose, succinate) and one shared among amino acids and nucleotides. Both coefficients were optimized via grid search to maximize agreement between predicted and measured growth curves across the entire dataset. The sugar scaler was searched over [0.001, 0.005, 0.01, 0.015, 0.02], and the amino-acid/nucleotide scaler over [0.1, 0.2, 0.3, 0.4, 0.5, 0.6, 0.7, 0.8, 0.9, 1.0]. The best-performing parameters, yielding the highest $R^2$, are 0.02 for sugars and 0.2 for amino acids/nucleotides. One notices from the results plotted in Fig. S9 that, as expected, dFBA does not handle lag-phase nor biomass depletion.

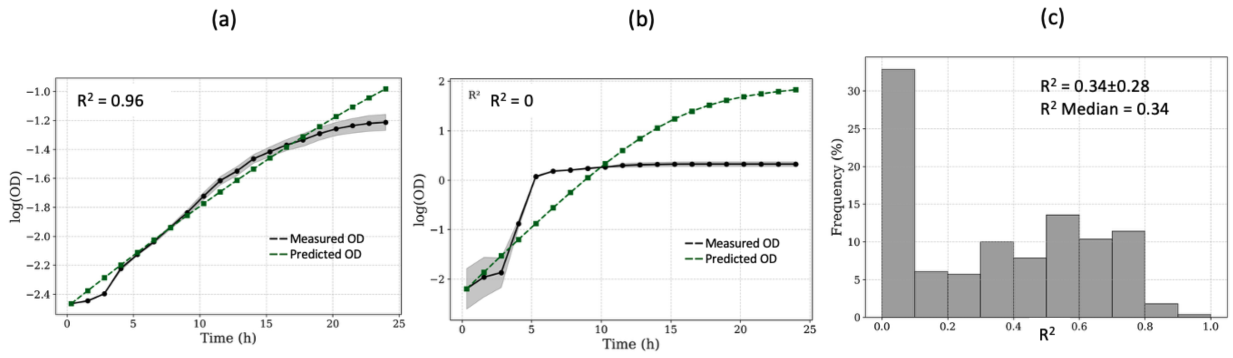


**Fig. S9. dFBA predicted growth curves for the *E. coli* M28 dataset.**

*(a) High $R^2$ example (medium number 276). (b) Low $R^2$ example (medium number 2). (c) Distribution of $R^2$ values between measured and dFBA predicted growth curves.*
